# Comparative analysis of the treatment-naïve microbiome across rheumatic diseases to predict MTX treatment response

**DOI:** 10.1101/2024.07.31.605913

**Authors:** Lena Amend, Kun D. Huang, Pavaret Sivapornnukul, Miriam Rabenow, Agata Anna Bielecka, Xiao-yu Wang, Meina Neumann-Schaal, Torsten Witte, Till Strowig

## Abstract

The human gut microbiota is recognized as a modulator of inflammatory diseases and has been linked to interindividual differences in therapy responsiveness. However, the robustness of disease-specific microbiome signatures across closely related diseases is rarely compared. Here, we compared treatment-naïve microbiota composition and functional potential across rheumatic diseases, including rheumatoid arthritis (RA) and spondyloarthritis subforms, to identify disease-specific biomarkers. While we failed to define robust disease-specific microbiota signatures, we identified microbial signatures linked to methotrexate (MTX) responsiveness for the two rheumatic diseases RA and psoriatic arthritis (PsA), for which MTX is the first-line treatment. Notably, the signatures were distinct, i.e., we could define a signature based on the relative abundance of microbial species for RA, yet the signature for PsA was based on the relative abundance of microbial pathways. Together this supports the previously recognized value of microbiota to predict treatment responses to MTX in RA and identifies distinct signatures predicting MTX responsiveness for PsA.

## Introduction

The human microbiome has been recognized over the last decade as a modifier of human diseases such as metabolic syndrome, inflammatory bowel disease, and colorectal cancer, contributing to disease initiation and progression by diverse mechanisms, for instance, the production of microbial metabolites [1–3] . More recently, the intestinal microbiome emerged also as an important factor regulating interindividual differences in responses to drugs by directly metabolizing compounds or by providing immunomodulatory signals. Prominent examples are the metabolism of digoxin, affecting its efficacy [4] , and checkpoint inhibitors, which display reduced activity in the absence of co-stimulating microbes [5] . Despite these advances, defining microbial signatures linked explicitly to disease initiation or treatment is frequently co-founded by environmental and genetic factors independently linked to disease as well as comorbidities and the intake of medications that directly modulate microbiota composition and activation [6].

Rheumatoid arthritis (RA) and spondyloarthritis (SpA) are two common rheumatic diseases driven by autoimmunity, but they differ in their causes, symptoms, and the parts of the body they typically affect. RA is characterized by joint inflammation, synovitis, and the presence of autoantibodies against citrullinated proteins (ACPA) and/or rheumatoid factor (RF) [7]. SpA comprises a group of inflammatory diseases that primarily affect the spine and sacroiliac joints but can also involve peripheral joints, entheses, and other organs. For instance, PsA, a subform of SpA, is often associated with heterogeneous manifestations on several body sites, including psoriasis, enthesitis, dactylitis, bursitis, peripheral arthritis, and axial symptoms [8,9].

Various microbiome signatures have been reported in treatment-naïve RA and SpA patients [10,11], yet, most studies focused on either of the diseases. Of note, despite the substantial differences within the underlying pathophysiology, the recommended first-line therapies for treating RA and PsA resemble each other. The European League Against Rheumatism (EULAR) recommendations advice orally administered low-dose Methotrexate (MTX) as the first treatment strategy in early RA patients [12]. Similarly, the prescription of MTX is recommended for new-onset PsA patients following an ineffective treatment with nonsteroidal anti-inflammatory drugs [13]. However, clinical studies evaluating MTX monotherapy efficacy consistently reported a substantial proportion of both patient groups not responding to medication. Specifically, remission rates of a maximum of 50 % and 67 % were found in MTX-treated RA and PsA patients, respectively [14–19]. Hence, the identification of factors responsible for clinical response and efficacy of MTX in rheumatic patients is of crucial importance to increase treatment success and improve disease progression. In addition, it would further allow the establishment of biomarkers to serve as prognostic indicators for MTX treatment response.

In RA patients, the factors of sex, age, symptom duration, and smoking status [20], as well as specific genetic polymorphisms [21], have been identified to influence the inter-individual variability in MTX response. In addition, the gut microbiota influences the metabolism and biotransformation of orally administered MTX, as specific groups of bacteria have been associated with detecting the MTX conversion products in the feces of rats [22]. More recently, the gut microbial composition of 26 treatment-naïve new-onset RA patients was found to differ between MTX responders and non-responders. Moreover, several bacterial gene orthologs and pathways were identified to be associated with clinical response to MTX [23]. In line with that, another study identified the composition of bacterial genes involved in MTX metabolism to differ between MTX responders and non-responders in 49 RA patients [24]. Together, these studies support the role of the microbiota in determining clinical response to MTX monotherapy in RA patients. However, the potential influence of the microbial community on MTX treatment efficacy has not yet been investigated in PsA patients.

In this study, we applied metagenomic sequencing to first compare the treatment-naïve microbiota across diverse rheumatic diseases, including RA and different forms of SpA (PsA, axial SpA (axSpA), reactive arthritis (ReA)), and a control group of individuals reporting similar rheumatic disease symptoms, yet with a diagnosis unrelated to rheumatic diseases. Next, we investigated the composition and metabolic capacity of the intestinal microbiota in treatment-naive newly diagnosed RA and PsA patients and its link to treatment response with orally administered MTX. The microbiota analysis across the analyzed rheumatic diseases identified a limited number of microbial species and pathways that differed in abundance when comparing individual groups but failed to define robust distinct disease-specific microbiota signatures. Nevertheless, we identified microbial signatures linked to MTX responsiveness for both RA and PsA. Notably, the signatures were distinct, i.e., we could define a signature based on the relative abundance of microbial species for RA, yet the signature for PsA was based on the relative abundance of microbial pathways, including the ability to produce short-chain fatty acids. Together this corroborates the previously recognized value of the gut microbiota to predict treatment responses to MTX in RA and identifies distinct signatures predicting MTX responsiveness for PsA.

## Results

### The new-onset gut microbiome of patients with different rheumatic diseases displays only minor differences in alpha diversity, composition, and function

The Rheuma-VOR cohort consists of patients with newly diagnosed rheumatic diseases and individuals diagnosed with non-rheumatic diseases (NRD). To compare the gut microbiome composition and functional potential of newly diagnosed patients with rheumatic diseases (RA, PsA, axSpA, and ReA) to the control group (NRD) and among each other, shotgun metagenomic sequencing of 239 fecal samples was performed (sequencing data size, mean = 2.4 Gb)(Table 1 and Figure S2). Next, species-level taxonomic and functional profiling was performed using MetaPhlan 4 and HUMAnN 3, respectively. The analysis of the taxonomic profiles did not identify significant differences in alpha diversity, i.e., within sample diversity, between the patient groups, except that the Shannon index was elevated in ReA patients compared to those with axSpA (Figure 1A, *P* = 0.05 Mann-Whitney U test). Similarly, beta diversity analysis of taxonomic profiles, i.e., between sample diversity, revealed no statistically significant differences associated with rheumatic disease types except for differential distribution of RA and axSpA samples on the second principal coordinate axis (PC2, Figure 1B, *P* = 0.05 Mann-Whitney U test). Similarly, the link between microbial metabolic pathways and rheumatic disease types appeared to be insignificant (P = 0.574, PERMANOVA test)

**Figure 1.**
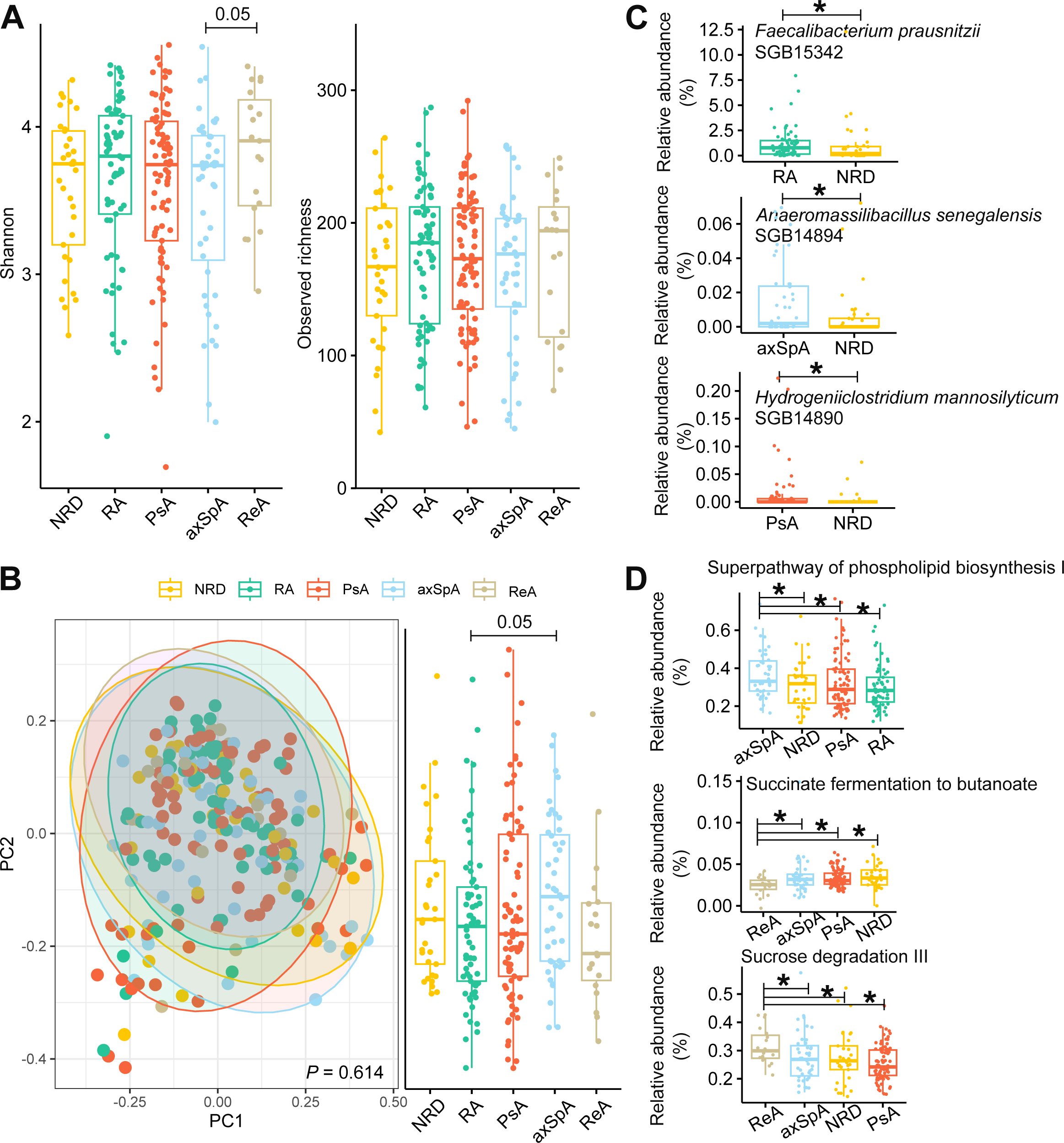
The structure of microbiome community composition in patients with NRD, RA, PsA, axSpA, and ReA. **A** Alpha diversity of samples from different disease groups. Insignificant differences in alpha diversity between groups were observed except for axSpA and ReA. **B** Beta diversity with samples colored according to disease groups (left side, PERMANOVA test). Samples from different disease groups were distributed on the second principal coordinate axis (right side, Mann-Whitney U test). **C** Example species differentially abundant between rheumatic diseases and NRD (see Supplementary Table 1 for a complete list). **D** Example metabolic pathways differentially abundant between rheumatic diseases and NRD (see Supplementary Table 1 for a complete list).

**Table 1:**
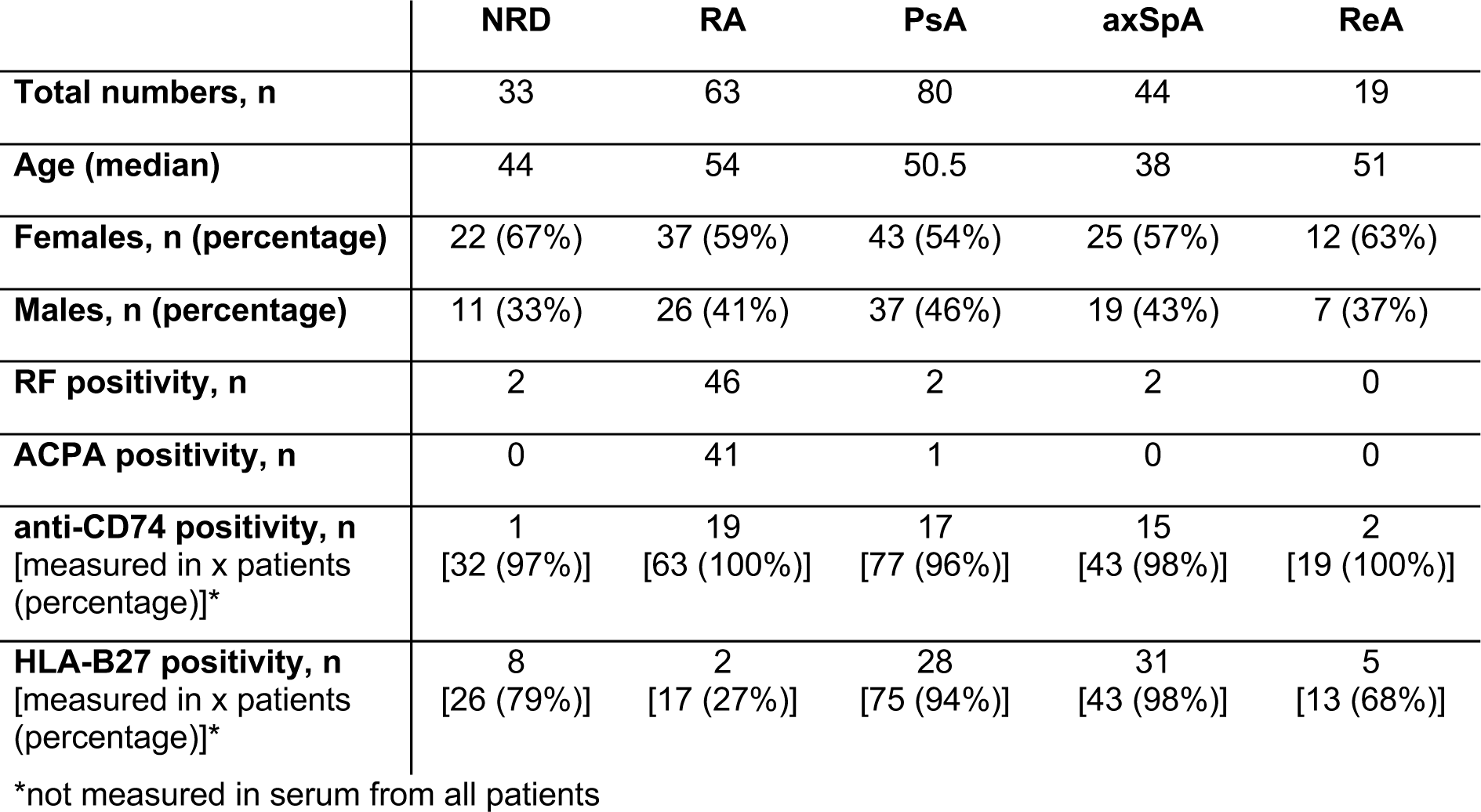
Overview of initial patient samples.

|  | **NRD** | **RA** | **PsA** | **axSpA** | **ReA** |
| --- | --- | --- | --- | --- | --- |
| **Total numbers, n** | 33 | 63 | 80 | 44 | 19 |
| **Age (median)** | 44 | 54 | 50.5 | 38 | 51 |
| **Females, n (percentage)** | 22 (67%) | 37 (59%) | 43 (54%) | 25 (57%) | 12 (63%) |
| **Males, n (percentage)** | 11 (33%) | 26 (41%) | 37 (46%) | 19 (43%) | 7 (37%) |
| **RF positivity, n** | 2 | 46 | 2 | 2 | 0 |
| **ACPA positivity, n** | 0 | 41 | 1 | 0 | 0 |
| **anti-CD74 positivity, n**<br>[measured in x patients<br>(percentage)]* | 1<br>[32 (97%)] | 19<br>[63 (100%)] | 17<br>[77 (96%)] | 15<br>[43 (98%)] | 2<br>[19 (100%)] |
| **HLA-B27 positivity, n**<br>[measured in x patients<br>(percentage)]* | 8<br>[26 (79%)] | 2<br>[17 (27%)] | 28<br>[75 (94%)] | 31<br>[43 (98%)] | 5<br>[13 (68%)] |

\*not measured in serum from all patients

To identify individual microbial species and metabolic pathways differentially abundant between rheumatic disease groups and NRD, a per-feature screening using MaAsLin 2 [25], which implements linear mixed models, was performed. We identified three species enriched in three rheumatic disease types compared to the control group (Figure 1C). First, *Faecalibacterium prausnitzii* (SGB15342), a known short-chain fatty acid (SCFA) producer, was more abundant in the samples of RA than in those of NRD (control). *Anaeromassilibacillus senegalensis* SGB14894, initially isolated from a kwashiorkor patient, displayed increased abundance in the gut microbiomes of patients with axSpA compared to the NRD group [26]. Interestingly, *Hydrogeniiclostridium mannosilyticum* SGB14890, which was reported to be associated with poorer response for immune checkpoint blockade treatment in melanoma [27], was enriched in the gut microbiome of PsA patients compared to NRD patients. Of note, nine species were also differentially abundant between rheumatic diseases, for example, the complex carbohydrate degrader *Ruminococcus bromii* SGB4285 was enriched in the microbiome of PsA patients compared to ReA patients (Supplementary Table 1). On the microbial functional level, we also identified 20 metabolic pathways enriched/depleted in one of the rheumatic disease groups compared to NRD and 16 differentially abundant pathways between two rheumatic disease groups (Supplementary Table 1). For instance, the abundance of the superpathway phospholipid biosynthesis I was considerably increased in axSpA patients compared to patients with NRD, PsA, and RA, respectively (Figure 1D). The pathway fermenting succinate to butanoate (butyrate) was significantly less abundant in ReA patients than in patients with NRD, axSpA, and PsA, respectively. On the contrary, the abundance of sucrose degradation III pathway was increased in ReA patients compared to those with NRD, axSpA, and PsA, respectively.

Together, this suggests the absence of distinctive microbiota signatures between untreated patients with different rheumatic diseases

### The taxonomic composition of the pre-treatment gut microbiome has predictive potential for MTX treatment outcomes for newly diagnosed RA patients

A previous study reported the association of pre-treatment gut microbiome to MTX treatment outcome in RA patients [23], but the predictive potential for other MTX-treated rheumatic diseases was not investigated. To characterize the impact of gut microbiota taxonomic composition on treatment success of orally administered MTX in the Rheuma-VOR cohort, PsA (n=28) and RA patients (n=29) undergoing MTX treatment following the rheumatic diagnosis were studied. Clinical response to MTX was determined by rheumatologists applying disease-specific rheumatic criteria (see methods), based on which patients were classified as MTX responders (MTX-R; PsA n=14, RA n=19) and MTX non-responders (MTX-NR; PsA n=14, RA n=10). Patient characteristics, including age, gender, symptom duration, smoking status, further PsA disease manifestations, autoantibody status, and medication intake before the initial rheumatic diagnosis were retrieved (Table 2). For both PsA and RA, MTX-R and MTX-NR patients were matched by metadata parameters, with only age and enthesitis manifestation significantly differing between MTX-R and MTX-NR for PsA (Table 2). The previously generated metagenomic dataset was reanalyzed for the PsA and RA patient subsets to compare the baseline microbiota structure between MTX-R and MTX-NR. Microbial alpha diversity analysis revealed an increased observed richness and Shannon index in MTX-R compared to MTX-NR for both PsA and RA patient groups, but the observed differences were not statistically significant (Figure 2A). The results for the RA patients contradict a previous study, which reported reduced bacterial diversity in MTX-R compared to MTX-NR [23].

**Figure 2.**
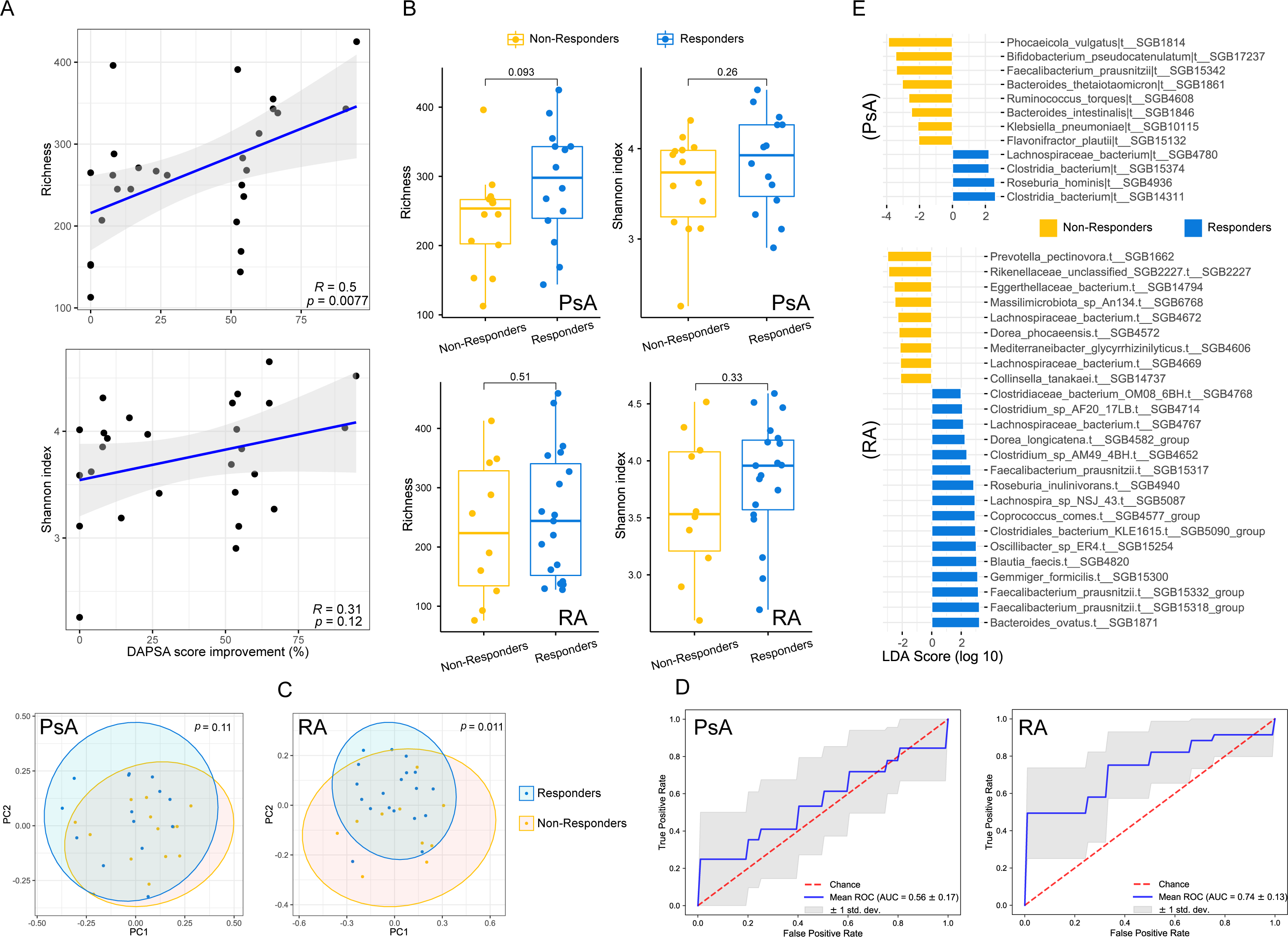
Taxonomic composition of the gut microbiome in PsA and RA patients and its predictive potential for treatment outcome. **A** Boxplot comparing the baseline microbiome richness (left side) and Shannon index (right side) in PsA (upper panel) and RA patients (lower panel) before MTX treatment initiation. **B** Correlation between DAPSA score improvement (before v.s. after MTX treatment) and richness (upper panel) and Shannon index (lower panel) in the PsA microbiome (Pearson’s correlation method). **C** Principal coordinate analysis (PCoA) based on microbial abundance profiled using MetaPhlAn4 between responders and non-responders in PsA (left panel) and RA (right panel). **D** ROC curve (Receiver operating characteristic curve) of the machine learning model based on the species-level relative abundances of responders and non-responders in PsA (left panel) and RA (right panel) using a random forest-based approach. **E** Species-level taxonomic biomarkers associated with responders and non-responders in PsA (upper panel) and RA (lower panel) were identified using LEfSe.

**Table 2:**
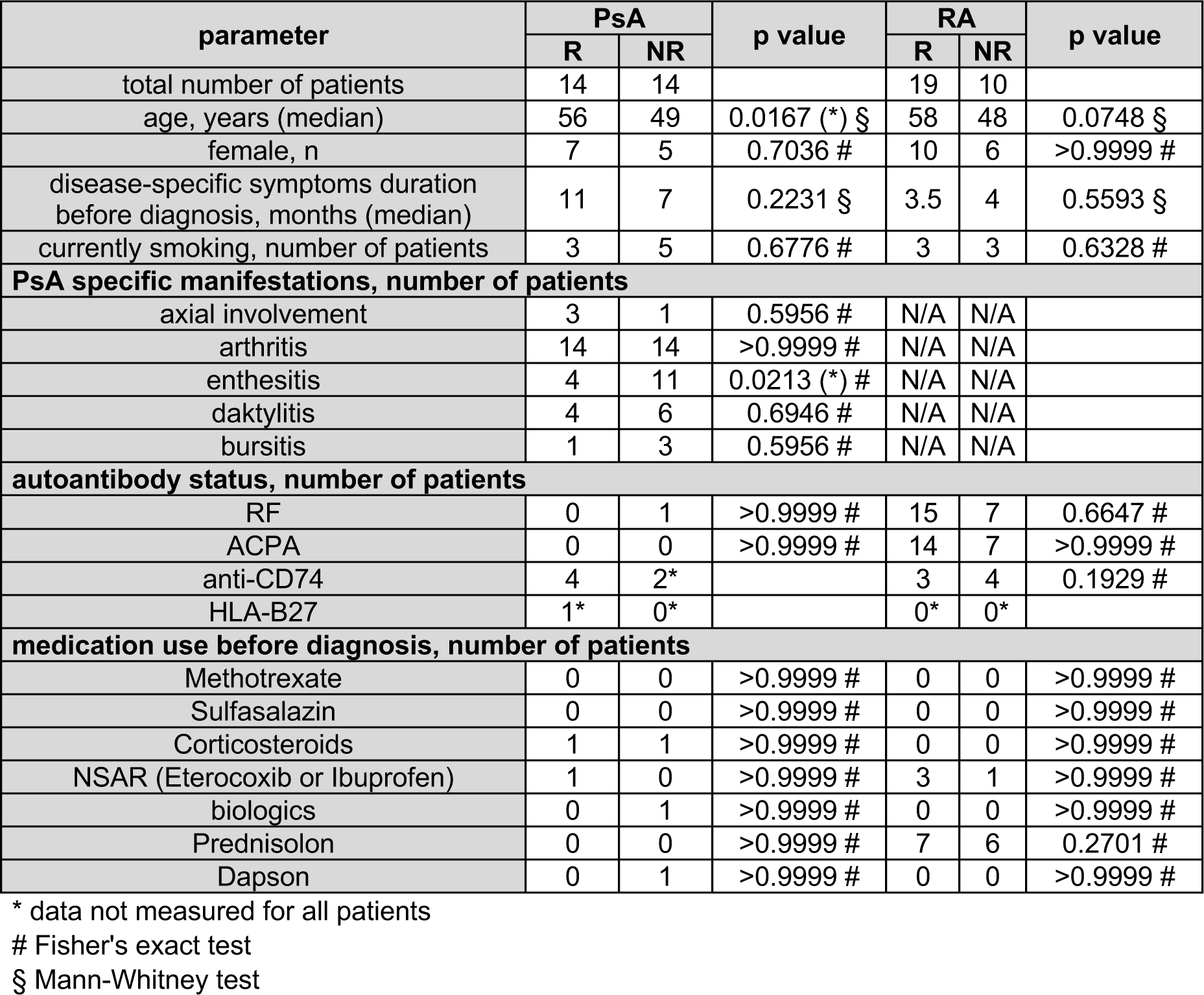
Overview metadata of MTX subcohort.

| parameter | PsA |  | p value | RA |  | p value |
| --- | --- | --- | --- | --- | --- | --- |
|  | R | NR |  | R | NR |  |
| total number of patients | 14 | 14 |  | 19 | 10 |  |
| age, years (median) | 56 | 49 | 0.0167 (*) § | 58 | 48 | 0.0748 § |
| female, n | 7 | 5 | 0.7036 # | 10 | 6 | >0.9999 # |
| disease-specific symptoms duration before diagnosis, months (median) | 11 | 7 | 0.2231 § | 3.5 | 4 | 0.5593 § |
| currently smoking, number of patients | 3 | 5 | 0.6776 # | 3 | 3 | 0.6328 # |
| **PsA specific manifestations, number of patients** |  |  |  |  |  |  |
| axial involvement | 3 | 1 | 0.5956 # | N/A | N/A |  |
| arthritis | 14 | 14 | >0.9999 # | N/A | N/A |  |
| enthesitis | 4 | 11 | 0.0213 (*) # | N/A | N/A |  |
| daktylitis | 4 | 6 | 0.6946 # | N/A | N/A |  |
| bursitis | 1 | 3 | 0.5956 # | N/A | N/A |  |
| **autoantibody status, number of patients** |  |  |  |  |  |  |
| RF | 0 | 1 | >0.9999 # | 15 | 7 | 0.6647 # |
| ACPA | 0 | 0 | >0.9999 # | 14 | 7 | >0.9999 # |
| anti-CD74 | 4 | 2* |  | 3 | 4 | 0.1929 # |
| HLA-B27 | 1* | 0* |  | 0* | 0* |  |
| **medication use before diagnosis, number of patients** |  |  |  |  |  |  |
| Methotrexate | 0 | 0 | >0.9999 # | 0 | 0 | >0.9999 # |
| Sulfasalazin | 0 | 0 | >0.9999 # | 0 | 0 | >0.9999 # |
| Corticosteroids | 1 | 1 | >0.9999 # | 0 | 0 | >0.9999 # |
| NSAR (Eterocoxib or Ibuprofen) | 1 | 0 | >0.9999 # | 3 | 1 | >0.9999 # |
| biologics | 0 | 1 | >0.9999 # | 0 | 0 | >0.9999 # |
| Prednisolon | 0 | 0 | >0.9999 # | 7 | 6 | 0.2701 # |
| Dapson | 0 | 1 | >0.9999 # | 0 | 0 | >0.9999 # |

\* data not measured for all patients

To enable a higher resolved analysis, microbial diversity and changes in disease activity were directly correlated. Notably, for PsA, a significant positive correlation was detected between observed richness (r=0.5, p=0.007 (**)) and DAPSA (disease activity in psoriatic arthritis) score improvement. This suggests that a diverse microbiota contributes to therapy success in PsA patients with orally administered MTX (Figure 2B). Given a wide variation in disease-specific symptom duration before rheumatic diagnosis between PsA patients (range), we repeated the analyses exclusively on PsA patients who had experienced symptom duration of i) less than one year (n= 20 patients) and i) less than 2 years (n= 23 patients)(five patients lacked relative data). Consistent with the results from PsA patients not sub-grouped by symptom duration, alpha diversity between MTX-R and MTX-NR did not differ significantly (Supplementary Figure 1C), while there was a significant, positive correlation between bacterial richness and disease activity in both symptom duration groups (Supplementary Figure 1B).

To investigate the taxonomic differences in the overall microbial community composition between MTX-R and MTX-NR, beta diversity was analyzed using Bray-Curtis dissimilarities and visualized by PCoA. In PsA patients, the overall microbial composition was not significantly different between MTX-R and MTX-NR (Figure 2C). In contrast, consistent with previous reports [23], beta diversity significantly differed between RA MTX-R and MTX-NR. Of note, the taxonomic microbial compositions of PsA and RA patients independent of MTX response did not differ from each other (Supplementary Figure 2C). Next, we used a Random Forest machine learning framework to estimate the prediction power of microbiota composition to differentiate MTX-R from MTX-NR. Only a weak microbiome-based prediction capability was observed for PsA (area under the receiver operating characteristic curve (AUC-ROC) 0.56), but a more profound performance to distinguish MTX-R and MTX-NR in between RA patients (AUC-ROC of 0.74) (Figure 2D). These results indicate that the gut microbiota taxonomic composition is potentially predictive of response to MTX in rheumatic patients, particularly in RA.

We further studied which individual microbial biomarkers are related to response to MTX therapy using linear discriminant analysis effect size (LEfSe) analysis (Figure 2E). In line with a more profound association between overall microbiome taxonomy and MTX response in RA (Figure 2C), we observed more bacterial species biomarkers related to MTX response in RA than PsA (Figure 2E). In PsA patients, four species were associated with MTX inefficiency, while eight were related to MTX response. MTX-NR in PsA harbored an increased abundance of three species belonging to the family *Bacteroidaceae (Phocaeicola vulgatus* SGB1814, *Bacteroides thetaiotaomicron* SGB1861, and *Bacteroides intestinalis* SGB1846) as well as increased abundances of the two health-associated bacterial species *Bifidobacterium pseudocatenulatum* SGB17237 and *Faecalibacterium prausnitzii* SGB15342. Furthermore, the MTX-NR in PsA patients had increased levels of *Klebsiella pneumonia* SGB10115, *Ruminococcus torques*

SGB4608, and *Flavonifractor plautii* SGB15132, which have all been linked to intestinal inflammation and diseases [28–30]. In contrast, the gut bacterial community of PsA patients responding to MTX treatment was enriched for species belonging to the families *Clostridiaceae* and *Lachnospiraceae*, commonly characterized by their SCFA-producing capacity [31–33].

LEfSe analysis in RA patients revealed stool samples of MTX-NR to harbor increased levels of nine distinct species, including species belonging to the family *Lachnospiraceace* (*Lachnospiraceae bacterium* SGB4672, *Dorea phocaeensis* SGB4572, *Lachnospiraceae bacterium* SGB4669, *Mediterraneibacter glycyrrhizinilyticus* SGB4606). Response to MTX in RA was characterized by 16 significantly enriched intestinal species, which included several species from the family *Clostridiaceae* and *Lachnospiraceae*. Additionally, *Faecalibacterium prausnitzii* (SGB15317, SGB15332, SGB15318), *Oscillibacter* sp SGB15254, and *Gemmiger formicilis* SGB15300 were enriched in RA MTX-R. These bacterial species belong to the *Oscillospiraceae*, a bacterial family implicated in butyrate production [34–36]. Thus, LEfSe analysis identified bacterial species associated with response to MTX in both patient groups and detected overlapping signatures (e.g. *Clostridiaceae* and *Lachnospiraceae*) linked to response to MTX in PsA and RA.

### Enrichment of microbial SCFA metabolic pathways in MTX responders with PsA compared to non-responders

Functional profiling using the HUMAnN3 pipeline [37] identified 548 and 569 pathways in PsA and RA patients included in the analysis of MTX treatment response, respectively. Response to MTX did not significantly impact the overall functional profiles in PsA nor in RA patients (Figure 3A). Random forest machine learning identified a moderate prediction capability of MTX response based on the functional potentials encoded in the gut microbiome in PsA patients (AUC-ROC 0.65), but not in RA patients (AUC-ROC 0.42) (Figure 3B). Similar to the taxonomic profiles, individual bacterial metabolic pathways associated with treatment response in PsA and RA patients were investigated by LEfSe analysis (Figure 3C). Interestingly, metabolic pathways appeared to share a stronger link with MTX response in the case of PsA, for which more individual biomarkers related to MTX response were identified compared to RA (n = 34 in PsA; n = 10 in RA). In PsA patients, 22 metabolic pathways were linked explicitly to MTX-R and 12 to MTX-NR. In both response groups, individual pathways related to amino acid biosynthesis (e.g. MTX-R: superpathway of sulfur amino acid biosynthesis: MTX-NR: L-valine biosynthesis) and carbohydrate metabolism (e.g. MTX-R: TCA cycle VI & I; MTX-NR: pentose phosphate pathway, superpathway of glucose and xylose degradation) were detected to be increased. Moreover, PsA MTX-R were specifically associated with pathways involved in SCFA metabolism (e.g., pyruvate fermentation to propanoate, glycerol degradation to butanol, pyruvate fermentation to butanol). In contrast, pathways involved in vitamin and cofactor metabolism (such as coenzyme A biosynthesis I & II and thiamine phosphate formation from pyrithiamine and oxythiamine) were enriched in MTX-NR. For RA, eight metabolic pathways were specifically associated with treatment inefficiency and two biomarkers with response (Figure 3C). MTX-NR displayed enrichment of pathways involved in fatty acid biosynthesis (such as phosphatidylglycerol biosynthesis I &II and lipid IVA biosynthesis) as well as amino acid metabolism (e.g., L-leucine degradation and L-isoleucine biosynthesis). At the same time, MTX-R were associated to the pathway CDP-diacylglycerol biosynthesis III and the superpathway of thiamine diphosphate biosynthesis II (respectively involved in fatty acid and vitamin biosynthesis). These results suggest that the functional capacity of intestinal microbes may contribute to inter-individual variation in MTX treatment outcomes, particularly for PsA patients.

**Figure 3.**
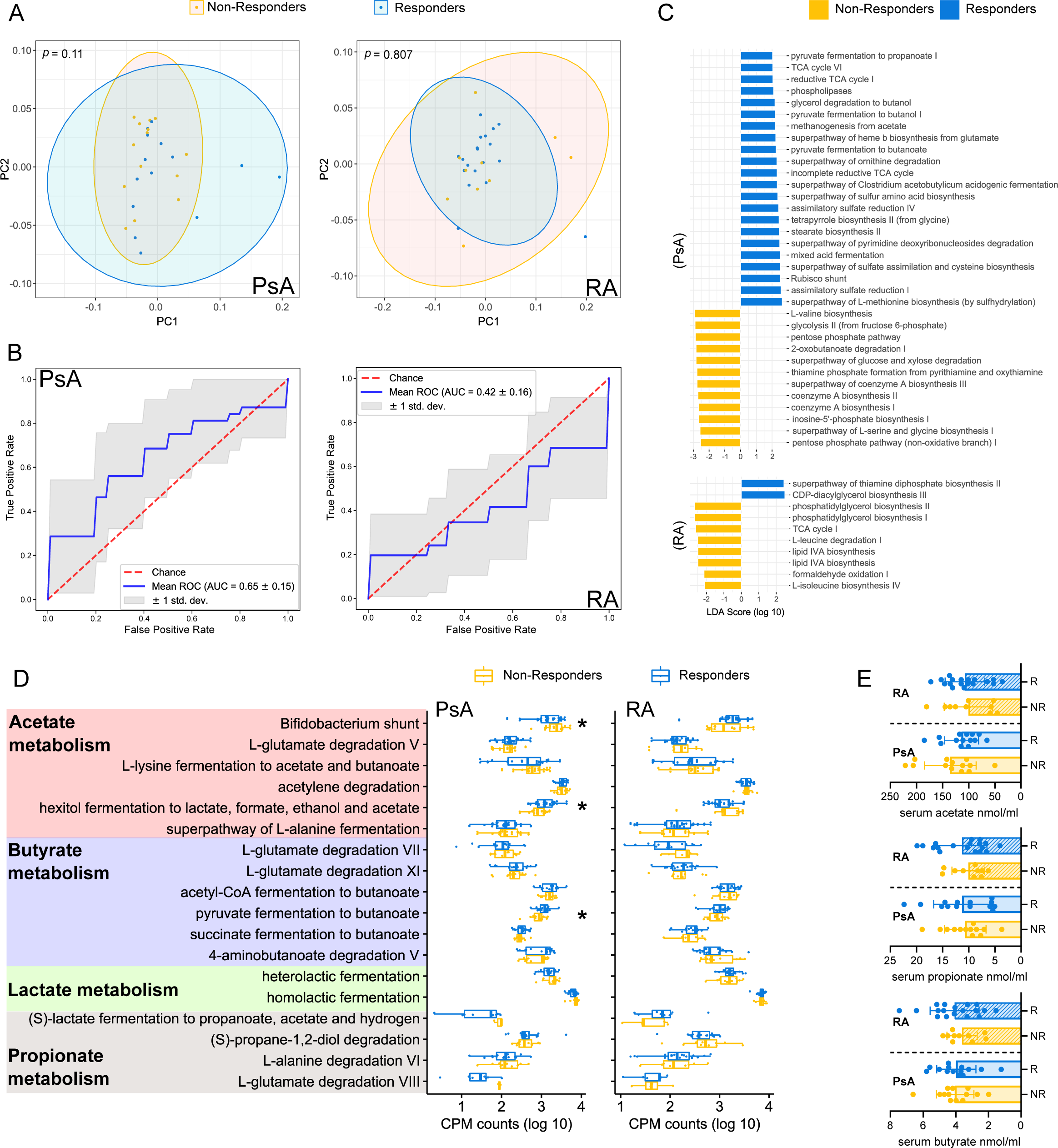
Metabolic pathways differentiating MTX responders and non-responders. **A** Principal coordinate analysis (PCoA) based on metabolic pathway abundances profiled using HUMAnN3 between responders and non-responders in PsA (left side) and RA (right side). **B** ROC curve (Receiver operating characteristic curve) of the machine learning model based on the HUMAnN3 metabolic pathway abundances of responders and non-responders in PsA (left side) and RA (right side) using a random forest-based approach. **C** Microbial metabolic pathway biomarkers associated with responders and non-responders in PsA (upper side) and RA (lower side) identified by LEfSe analysis. **D** Log abundance (copies per million) of pathways involved in SCFA production in responders and non-responders PsA (left side) and RA patients (right side). **E** Acetate, propionate, and butyrate concentration in the serum of non-responders (n=13) and responders (n=14) in PsA patients as well as non-responders (n=10) and responders (n=18) in RA patients.

As LEfSe analysis highlighted SCFA metabolism associated with MTX therapy success in PsA patients, we specifically quantified the levels of 16 bacterial pathways involved in SCFA metabolism in MTX response for PsA and RA patients (Figure 3D). In line with LEfSe analysis, SCFA pathway levels did not differ between MTX-R and MTX-NR in RA patients. In PsA patients, two SCFA pathways (“hexitol fermentation to lactate, formate, ethanol, and acetate” and “pyruvate fermentation to butanoate”) were increased in MTX-R and suggested an enhanced bacterial SCFA metabolism be linked to clinical response to MTX. Interestingly, the pathway “Bifidobacterium shunt”, a unique *Bifidobacterium* pathway producing acetate, was increased in PsA MTX-NR and corresponded to the detection of enriched *Bifidobacteria* species in MTX-NR by LEfSe. To identify a potential connection between SCFA pathway prediction and serum levels of SCFAs in the patients, acetate, butyrate, and propionate concentrations were quantified using targeted metabolomics in the serum of PsA and RA patients clustered by response to MTX. Neither in PsA nor RA, concentrations of the three SCFAs differed significantly between MTX response groups (Figure 3E). Hence, serum SCFA concentration did not reflect the SCFA pathway prediction based on metagenomics sequencing.

## Discussion

Rheumatic diseases have been associated with alterations in the microbiome composition by several studies in the past decade [10,38,39]. For instance, an increased prevalence and relative abundance of bacteria belonging to the genus Prevotella (recently reclassified into the genus Segatella) were reported to be associated with new-onset RA [40]. However, inconsistencies between study outcomes have been noted [11], potentially originating from variations in cohort-based factors, e.g., disease duration and activity, drug intake, and geographic origin, as well as from differences in methodologies, including sampling protocols, sequencing methods, and data analysis. In this study, we analyzed samples from patients recruited through the German Rheuma-VOR cohort, which enrolls patients after referral by their primary care physician and before a final diagnosis is determined and thus therapy started. This enabled us to analyze the gut microbiome of new-onset, treatment-naïve rheumatic patients from the same geographic origin, thereby minimizing the effect of confounding factors on the gut microbiome composition. The control group in our study consisted of patients with symptoms of back or joint pain but with a negative diagnosis for any rheumatic disease, which is distinct from other studies recruiting healthy donors as controls. Interestingly, we found that the gut microbiome diversity and its global structure did not differ between rheumatic diseases and the controls during disease onset. Moreover, only a few bacterial species were identified to be differentially abundant when comparing the groups. Of note is that members of the Prevotellaceae family did not display differences in abundance or prevalence across the different rheumatic diseases. Hence, our data suggests that the intestinal microbiota is not significantly perturbed within the timeframe of rheumatic disease onset but might undergo alterations later on during disease progression and therapy initiation.

Orally administered MTX is a standard first-line medication for new-onset PsA and RA patients [12,13], but there is a high variability in the treatment efficacy among individuals [14–19]. Recent studies suggested that the gut microbiome diversity, composition, and functional potential are associated with response to MTX in RA patients [23,24]. In this study, no associations between microbiome diversity or global composition and MTX response were observed for new-onset PsA and RA patients. However, the enrichment of bacterial species in the RA MTX response group, and to a more limited degree in the PsA MTX response group, identified by LEfSe, suggests specific bacteria to be linked to clinical response. The consistently increased abundance of the SCFA-producing bacterial families *Lachnospiraceae* and *Clostridiaceae* among PsA and RA responders, together with the enrichment of SCFA production pathways in PsA MTX-R based on functional prediction, associates SCFAs with MTX therapy success. Of note, the increase in SCFA metabolism pathways in PsA MTX-R does not necessarily imply SCFAs to enhance MTX response. Instead, the enrichment of SCFA pathways could merely be a marker for the prevalence of SCFA-producing microbes, which beneficially affect MTX treatment independently of SCFA production. Hence, future investigations are needed to determine the impact of SCFAs and SCFA-producing bacteria on MTX efficacy in rheumatic diseases. Nevertheless, SCFAs have previously been identified to positively affect the treatment-independent pathogenesis of rheumatic diseases [41,42]. Moreover, a recent study identified butyrate supplementation to inhibit MTX-induced gastrointestinal mucositis in an organoid model, potentially indicating that butyrate minimizes MTX’s cell toxicity and manipulates MTX uptake [43]. In contrast to the observed enrichment of SCFA-producing bacteria and SCFA pathways in MTX-R, serum SCFA levels did not differ among response groups. It needs to be noted, however, that acetate, propionate, and butyrate are heavily metabolized by colonocytes and hepatocytes upon their production, resulting in only a minor fraction eventually reaching the systemic circulation [44]. Hence, the quantification of SCFAs in the serum in patients does not reflect actual SCFA levels in the intestine and can, therefore, only be interpreted cautiously.

Collectively, these insights highlight the intestinal microbiota as a crucial target for enhancing the therapeutic efficacy of the indispensable cDMARD MTX in two rheumatic diseases.

## Methods

### Cohort description

The Rheuma-VOR cohort is an ongoing, prospective German cohort comprising individuals with symptoms of rheumatic diseases. Patients fulfilling the Berlin criteria or showing arthritis-specific symptoms (swollen joints, morning stiffness for > 30 min with or without elevated C-reactive protein (CRP) or erythrocyte sedimentation rate (ESR)) were referred from their general practitioner to rheumatologists. PsA and RA were diagnosed using screening questionnaires based on ACR classification criteria for RA and the CASPAR classification criteria, as well as the PEST and EARP questionnaires for PsA. Patients with nonspecific symptoms (such as back pain and/or arthralgia) who were neither diagnosed with RA or PsA nor with other rheumatic diseases were classified as NRD patients. Here, we reported a total of 239 patients (Figure S2).

MTX treatment success was assessed based on the EULAR response criteria for RA and the DAPSA score for PsA (Table 2). For RA, “responders” were defined as patients who responded well to MTX, while patients with no response or moderate response were classified as “non-responders”. For PsA, the term “responders” included patients whose DAPSA scores decreased by 50% compared to baseline, while patients with higher DAPSA scores were classified as “non-responders”.

### Sample collection, DNA extraction, and metagenomic sequencing

Native stool samples were collected in Quick separ tubes (prefilled with RNA separ, Cat: 6480 RNA), shipped to the laboratory at room temperature, and frozen upon arrival at -80°C until DNA isolation. DNA was extracted using the ZymoBIOMICS 96 MagBead DNA Kit following the manufacturer’s instructions. Metagenomic libraries were prepared using the Illumina DNA PCR-Free Library Kit and the IDT for Illumina DNA/RNA UD Indexes. Sequencing was performed on the NovaSeq S4 PE150 platform with a 25 Mio reads/sample sequencing depth. Metagenomic reads were quality-checked and processed using BBMap (sourceforge.net/projects/bbmap/) with the Ensembl masked human genome GRCh38 and phiX.

The species-level microbial community was profiled on pre-processed reads by running MetaPhlAn 4.0 [47] using default settings. The analysis did not consider species with a minimum relative abundance < 0.0001. Microbial abundance profiles were merged using merge_metaphlan_tables.py, a utility script in MetaPhlAn 4.0. HUMAnAN 3.0 [37] was applied to profiling microbial metabolic pathways on metagenomic reads with default settings. The Python script humann_renorm_table.py, as part of the pipeline, was used to normalize metabolic pathways to relative abundance, and individual profiles were subsequently converted into a merged abundance table using the utility script humann_join_tables.py.

### Calculation of alpha and beta diversity

Alpha diversity was investigated by calculating observed richness and Shannon index by R package *mia* (https://github.com/microbiome/mia) based on species-level relative abundances. Beta diversities were calculated using Bray-Curtis dissimilarity based on relative abundances of both profiled species and metabolic pathways, followed by a principal coordinates analysis (PCoA) using the R package *ggpubr* v0.4.0 (https://rpkgs.datanovia.com/ggpubr). A statistical significance test of beta diversities was conducted using a PERMANOVA analysis using the R package *vegan* v2.6.2 [48].

### Taxonomic and functional biomarker analysis

To identify species and metabolic pathways differentially abundant between rheumatic diseases, we first performed linear mixed models implemented in a biomarker identification tool MaAsLin 2 [25], setting as fixed effects disease type, age, and gender and using logarithm transformation. To identify biomarkers association with responders or non-responders to MTX treatment, we used a similar method - which is more sensitive for small datasets - linear discriminant analysis effect size (LEfSe) with default parameter settings [49]. Features with a logarithmic LDA score over 2.0 were considered significantly discriminant.

### Random forest-based machine learning analysis

Machine learning experiments in this study were based on the random forest model implemented by a python package scikit-learn v1.1.2 [50,51] as it has been shown to outperform other approaches in training microbiome data [52,53]. To create a learning model, we set 1,000 estimator trees and Shannon entropy to evaluate the quality of node splitting of a tree. As suggested elsewhere [54,55], we assigned at least one sample per leaf and 30% of features for a tree in the hyperparameter setting. We fit the model to the dataset comprising responders and non-responders from RA and PsA, respectively. The prediction power for each dataset was evaluated by stratified 5-fold cross-validation to ensure enough balanced binary cases in each partition. We repeated the procedure of dataset folding and model evaluation 20 times with a result of an average of over 100 validation folds per practice.

### Prediction of short chain fatty acid metabolism in intestinal microbiota

To investigate the SCFA metabolism potential between two groups, 28 MetaCyc-archived metabolic pathways involved in the SCFA fermentation [56] were extracted (see below), and abundances were calculated based on HUMAnAN 3.0 outputs.

| **Pathway** | **Class** |
| --- | --- |
| acetate and ATP formation from acetyl-CoA I | Fermentation to Acetate |
| acetate and ATP formation from acetyl-CoA II | Fermentation to Acetate |
| acetate and ATP formation from acetyl-CoA III | Fermentation to Acetate |
| acetate formation from acetyl-CoA (succinate) | Fermentation to Acetate |
| acetylene degradation (anaerobic) | Fermentation to Acetate |
| Bifidobacterium shunt | Fermentation to Acetate |
| glycolate degradation II | Fermentation to Acetate |
| hexitol fermentation to lactate, formate, ethanol and acetate | Fermentation to Acetate |
| L-glutamate degradation V (via hydroxyglutarate) | Fermentation to Acetate |
| L-lysine fermentation to acetate and butanoate | Fermentation to Acetate |
| superpathway of L-alanine fermentation (Stickland reaction) | Fermentation to Acetate |
| 4-aminobutanoate degradation V | Fermentation to Butyrate |
| acetyl-CoA fermentation to butanoate | Fermentation to Butyrate |
| L-glutamate degradation V (via hydroxyglutarate) | Fermentation to Butyrate |
| L-glutamate degradation VII (to butanoate) | Fermentation to Butyrate |
| L-glutamate degradation XI (reductive Stickland reaction) | Fermentation to Butyrate |
| L-lysine fermentation to acetate and butanoate | Fermentation to Butyrate |
| pyruvate fermentation to butanoate | Fermentation to Butyrate |
| succinate fermentation to butanoate | Fermentation to Butyrate |
| Bifidobacterium shunt | Fermentation to Lactate |
| heterolactic fermentation | Fermentation to Lactate |
| hexitol fermentation to lactate, formate, ethanol and acetate | Fermentation to Lactate |
| homolactic fermentation | Fermentation to Lactate |
| (S)-lactate fermentation to propanoate, acetate and hydrogen | Fermentation to Propionate |
| (S)-propane-1,2-diol degradation | Fermentation to Propionate |
| L-alanine degradation VI (reductive Stickland reaction) | Fermentation to Propionate |
| L-glutamate degradation VIII (to propanoate) | Fermentation to Propionate |
| superpathway of L-alanine fermentation (Stickland reaction) | Fermentation to Propionate |

### SCFA measurements in human serum samples

Acetate, butyrate, and propionate levels were measured in the patient’s serum samples stored at -80°C. Up to 80 µl of the serum was filled up with water until 400 µl and was further mixed with 600 µl H_2_O/internal standard (o-cresol), 60 µl of H_2_SO_4_ (ms grade), and 200 µl methyl tert-butyl ether (MTBE). Mixtures were vortexed for 5 min and centrifuged for 5 min at maximum speed. The ether phase was analyzed by GC-MS as previously described [57], using an Agilent GC-MSD system (7890B coupled to a 5977 GC) equipped with a high-efficiency source (HES).

### Statistical analysis

Statistical significance between groups was assessed using the Mann-Whitney U test. Statistical significance for multivariate analysis was performed using the PERMANOVA test. Correlation analysis was performed using the Pearson method. All analyses were conducted in the R v4.1.0 environment [58].

