## Supplementary Information for "Comparative analysis of the treatment-naïve microbiome across rheumatic diseases to predict MTX treatment response"

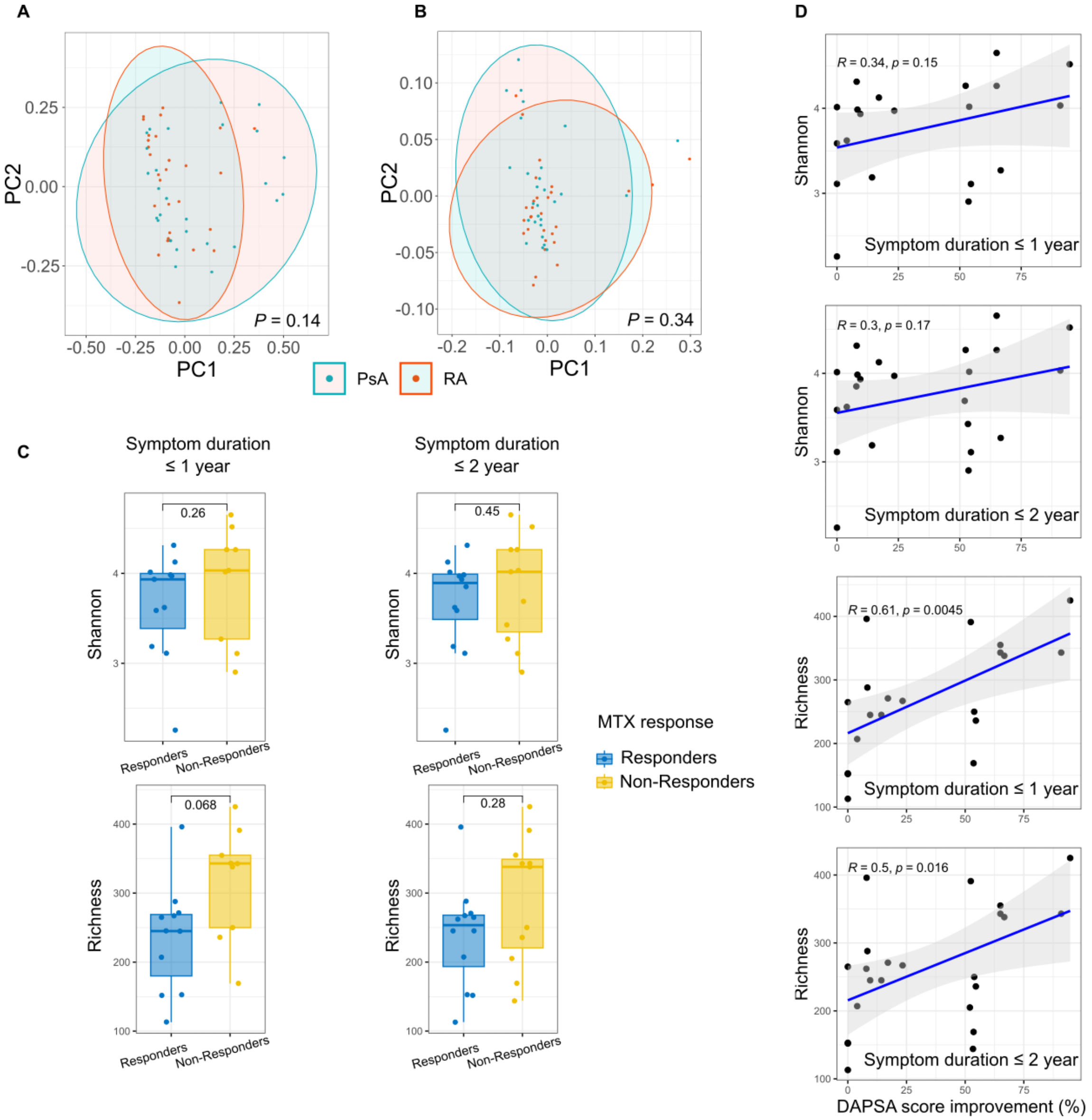


**Figure S1. A** and **B** PCoA plots based on taxonomic microbial compositions (C) and metabolic pathway abundance (D) including all PsA and RA patients non-clustered by MTX response. **C** Shannon index (upper panel) and microbial richness (lower panel) in PsA MTX-R and MTX-NR including all patients (left panels), only patients with symptom duration < 1 year (middle panels) and only patients with symptom duration < 2 years (right panels) prior to MTX treatment. **D** Correlation between Shannon index or microbial richness with DAPSA score improvement in PsA patients with symptom duration < 1 year (panel 1 and 3) and patients with symptom duration < 2 years (panel 2 and 4).

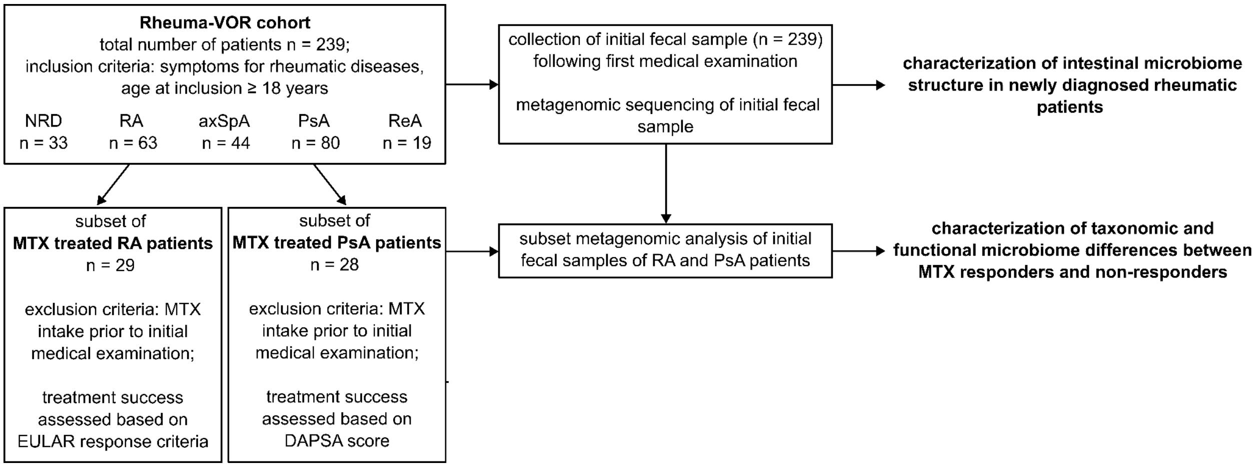


**Figure S2. Patient selection criteria for the whole Rheuma-VOR cohort and the subsets.** The analysis based on the whole Rheuma-VOR cohort included only initial samples from newly diagnosed patients with five groups of rheumatic diseases. The subsets included only RA and PsA patients receiving MTX treatment.

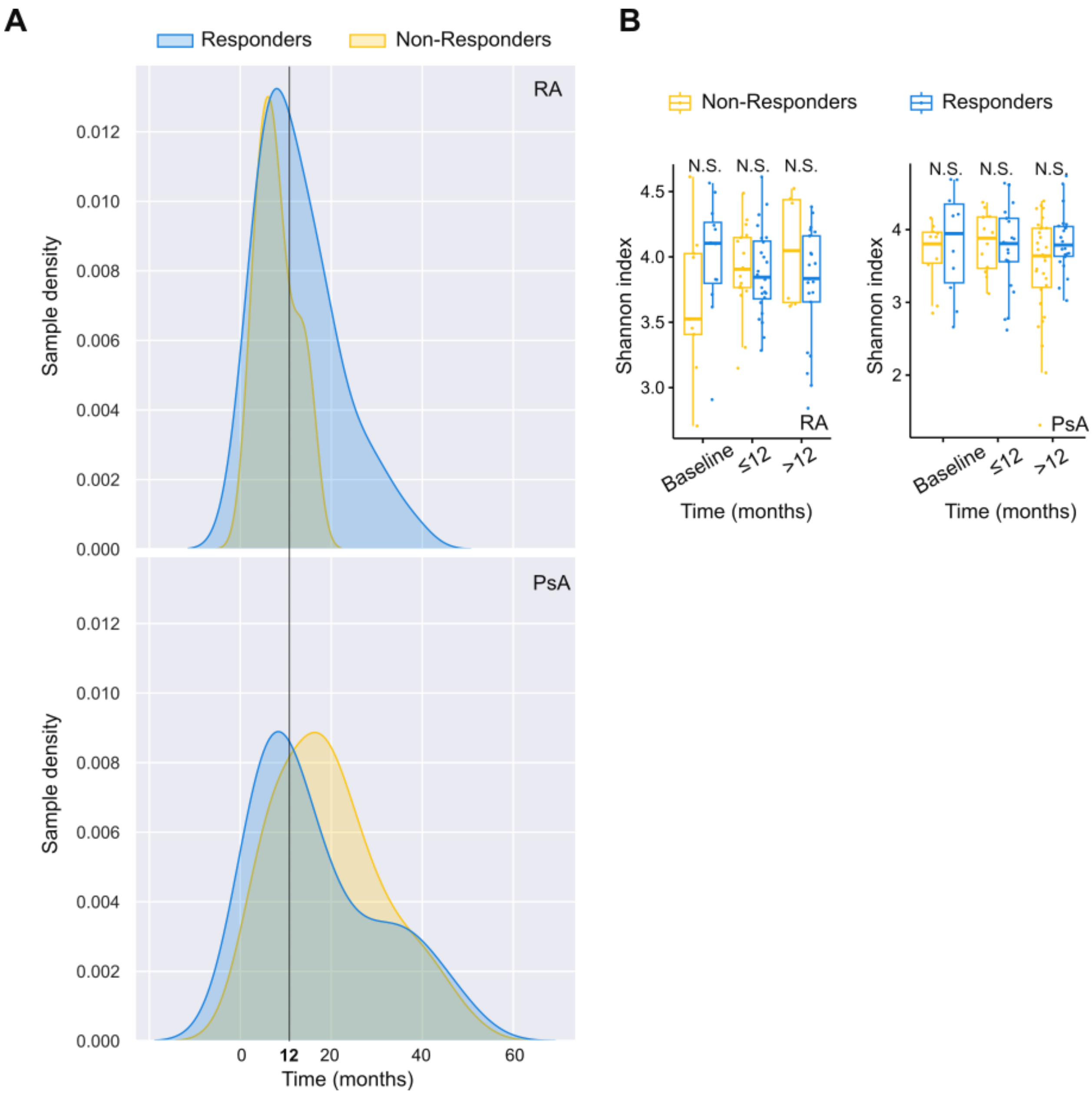


**Figure S3. A** The distribution of sample collection times in responders and non-responders of both RA and PsA patients. **B** Comparisons of the gut microbiome Shannon index between responders and non-responders based on samples collected at baseline, ≤12 and >12 months.

**Supplementary Table 1**: Microbial species or metabolic pathways differentially abundant between two groups of rheumatic diseases (q-value < 0.25).
